# ANGPTL3 binds to PCSK9 to coordinately regulate intracellular lipid homeostasis

**DOI:** 10.1101/2025.01.05.631413

**Authors:** Simone Bini, Valeria Pecce, Alessandra Pinzon Grimaldos, Alessia Di Costanzo, Ilenia Minicocci, Stella Covino, Daniele Tramontano, Silvia Piconese, Laura D’Erasmo, Marcello Arca

## Abstract

ANGPTL3 and PCSK9 are core proteins involved in lipid metabolism. Literature evidence highlights a potential mutual regulation between them: patients harboring loss-of-function variants in ANGPTL3 show reduced levels of circulating PCSK9. After predicting their interaction using an in-silico model, we verified the direct interaction between ANGPTL3 and PCSK9 through co-immunoprecipitation. The ANGPTL3-PCSK9 complex persisted under *fasting* conditions and dissociated under *feeding* conditions. Treatment with human LDLs was sufficient to simulate the feeding response. Then, we observed that the overexpression of PCSK9 enhances the uptake of LDLs that are not further metabolized, while the overexpression of ANGPTL3 enhances LDL turnover. The overexpression of both proteins restored LDL uptake and degradation, which became comparable to those of control cells. In conclusion, our findings indicate the existence of an ANGPTL3-PCSK9 complex, which coordinately contributes to the regulation of intracellular lipid homeostasis, aiming to prevent cellular metabolic overload.

**GRAPHICAL ABSTRACT:** Graphical abstract
Mechanistic model of ANGPTL3-PCSK9 complex function in the regulation of lipoprotein metabolism
Increased uptake of extracellular nutrients determines the separation of the ANGPTL3-PCSK9 complex. The two free proteins have different functions (1). Increased levels of PCSK9 determine an increased intracellular entrapment of circulating LDLs (2) that are not catabolized unless there is a presence of high intracellular levels of ANGPTL3. Increased ANGPTL3 levels also favor ApoB lipidation and secretion (3), whereas an increase in intracellular PCSK9 increases intracellular ApoB degradation. The increase in intracellular PCSK9 also determines a block in lipogenesis and favors beta-oxidation, thus decreasing the intracellular lipid content (4).

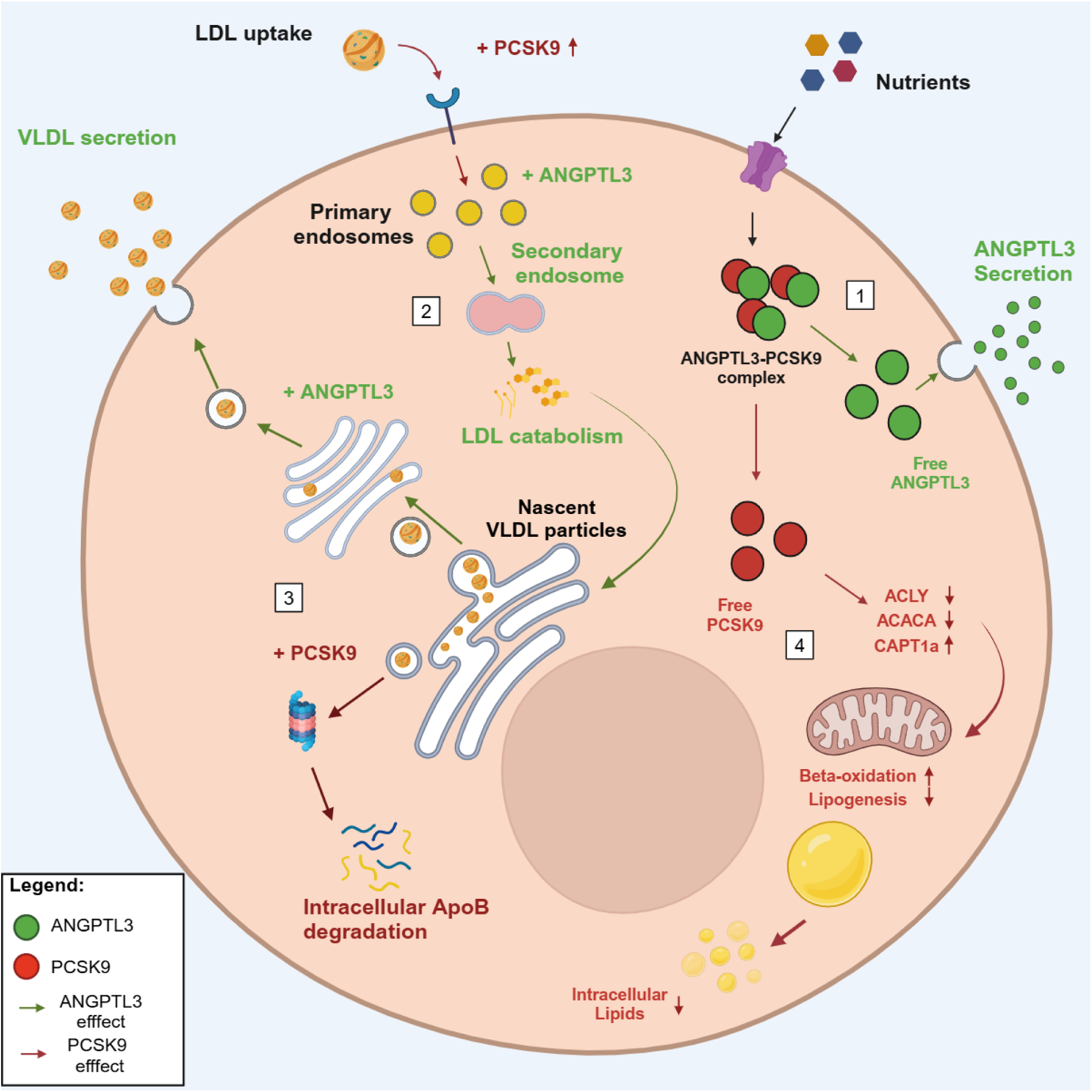

## INTRODUCTION

Angiopoietin-like protein 3 (ANGPTL3) and proprotein convertase subtilisin/kexin-like 9 (PCSK9) are known to be both essential regulators of plasma lipoprotein trafficking ^1–3^.

In circulation, ANGPTL3 inhibits the activity of extracellular lipases, particularly lipoprotein lipase and endothelial lipases ^4^. Through these actions, ANGPTL3 plays a pivotal role in the removal of triglyceride-rich lipoproteins (TRLs), very-low-density lipoproteins (VLDL), and chylomicrons, as well as in the catabolism of high-density lipoproteins (HDL) ^1,5^, in particular, through the regulation of TRLs lipolysis in the fasting and postprandial phase, ANGPTL3 has also been attributed a role in the distribution of lipid energy substrates between storage and highly metabolizing tissues^6^. These effects in humans have been firmly established by investigating individuals harboring loss-of-function variants (*LOF*) in the *ANGPTL3* gene. They show a comprehensive reduction of plasma concentration of TRLs, HDL, and low-density lipoprotein cholesterol (LDL-C), a phenotype named Familial Hypobetalipoproteinemia type 2 (FHBL2)^7^. FHBL2 is an ultra-rare condition, and affected individuals appear to be protected from cardiovascular disease and do not show an increased prevalence of liver steatosis^8–13^

The *PCSK9* belongs to the superfamily of kexin-like proteases, but its proteolytic activity is lost during the protein maturation. Indeed, the PCSK9 pro-peptide is self-cleaved in the endoplasmic reticulum, folds separately from the rest of the protein, and sets in its catalytic site, inactivating the complex^14,15^. The main action of PCSK9 is to regulate the plasma concentration of LDL-C by controlling the recycling of the low-density lipoprotein receptor (LDLR). PCSK9 can form a complex with LDLR, probably extracellularly, thus regulating LDLR turnover and its recycling on the cell membrane^16,17^.

It is currently thought that these two proteins may act in separate cellular compartments, influencing different lipids and lipoprotein pathways. Surprisingly, *Fazio et al.*^18^ found reduced levels of circulating PCSK9 in patients carrying homozygous LOF in ANGPTL3, thus implying a certain level of mutual interaction between ANGPTL3 and PCSK9. Additionally, *Tromp et al.*^19^ have recently observed that HDL and LDL carry a fraction of ANGPTL3, while PCSK9 is known to be carried by LDLs^14^. However, the possibility that ANGPTL3 and PCSK9 may interact with each other and undergo coordinated regulation at intracellular levels has never been considered.

Based on the evidence above, this work investigated the relationship between ANGPTL3 and PCSK9 regarding possible direct interaction and coordinated effects on lipid intracellular metabolism. The experiments were carried out in hepatic cell lines overexpressing *ANGPTL3*, *PCSK9,* or both genes to enhance protein activity over the background and in the context of variable nutrient availability, as it has been reported that both ANGPTL3 and PCSK9 proteins are sensitive to nutritional status^20,21^.

## MATERIALS AND METHODS

### In silico prediction of complex formation

We used AphaFold2 software to verify the possible interaction between ANGPTL3 and PCSK9 and to model a potential molecular docking into a bigger complex. AlphaFold2 produces a per-residue confidence metric called the predicted local distance difference test (pLDDT) on a scale from 0 to 100^22^. The pLDDT estimates how well the prediction would agree with an experimental structure based on the local distance difference test Cα (lDDT-Cα)^23^. The program computes the potential existence of the spatial distribution of the single residue sidechains in association with the potential position of the nearby residues based on existing crystallographic reposited in in the Protein Data Bank (PDB)^22^. A pLDDT > 90 is considered very accurate, pLDDT ≥ 70 is considered accurate^22^. Deriving from single residues pLDDTs the software computes the predicted template model score (pTM), this is a metric for the similarity between protein structures (in this case, between the predicted structure and the assumed real structure), the model is considered accurate if pTM > 0.50; for a pTM = 0.50 the *P*-value for the existence of the model is 5.5 × 10^−7^ ^24^. Similarly to pTM, the interface predicted model score (ipTM) computes the potential existence of the single residue sidechain position in comparison to the interfacing one, considering true the position of the remaining sidechains, like the pTM, also a structure having ipTM > 0.50 is considered accurate^25^.

### Cell line culture and transfection

HepG2, Hela and HEK-293 cell lines (provided by ATCC) were cultured in Dulbecco’s Modified Eagle Medium (Gibco-BRL Division, Thermo Fisher Scientific, Waltham, Massachusetts, USA) containing 10% fetal bovine serum (Gibco-BRL Division, Thermo Fisher Scientific, Waltham, Massachusetts, USA) and antibiotic–antimycotic solution (Gibco-BRL Division, Thermo Fisher Scientific, Waltham, Massachusetts, USA) and incubated at 37 °C in an atmosphere of 5% CO2. We generated the *ANGPTL3* and *PCSK9* overexpressing lines using plasmids containing the complementary DNA (cDNA) of *ANGPTL3* (NM_014495) or *PCSK9* (NM_174936) under the control of cytomegalovirus constitutive promoter (CMV). OriGene provided both pCMV-ANGPTL3 and pCMV-PCKS9 plasmids. Cell lines were transfected with empty vector or pCMV-ANGPTL3, pCMV-PCSK9, or both plasmids using Lipofectamine 3000 (Invitrogen Division, Thermo Fisher Scientific, Waltham, Massachusetts, USA) according to the manufacturer’s instructions. Cells were plated the day before transfection at 70% of confluent in a 6-well plate and starved for two hours before transfection. Cells were starved either in Dulbecco’s Modified Eagle Medium (Gibco-BRL Division, Thermo Fisher Scientific, Waltham, Massachusetts, USA) to simulate a nutrient-rich condition, and Ham’s F12 (Gibco-BRL Division, Thermo Fisher Scientific, Waltham, Massachusetts, USA) to simulate a nutrient-poor condition. The switch between nutrient-poor and nutrient-rich conditions was used to simulate *feeding*; conversely, switching from nutrient-rich to nutrient-poor medium was used to simulate *fasting*. Cells were harvested 48h after the transfections.

### Protein extraction

Total proteins were extracted from cell pellets with a lysis buffer containing Tris-HCl (pH 7.4, 50 mM), NaCl (150 mM), Triton (1% v/v), ethylenediaminetetraacetic acid (EDTA, 20 mM), phenylmethylsulfonyl fluoride (PMSF, 2 mM), protease and phosphatase inhibitors (Pierce, Rockford, IL, USA), leupeptin (2 ug/mL), glycerol (10% v/v). All extractions were done on ice, and the products were quantified using the Bradford method described by Pecce et al. in 2020 ^26^. Proteins from cell culture media were concentrated using StrataClean resin (Agilent). In particular, 2 mL of culture medium were incubated overnight with 10 uL of StrataClean resin in low-speed wheel rotation at 4°C. After overnight incubation, samples were centrifuged at 4000 RPM for 4 minutes, the supernatant was discharged, and the resin pellet containing proteins was suspended in 20 uL of loading buffer and incubated for 5 minutes at 95°C. Then, the samples were centrifuged at 4000 RPM for 4 minutes, and supernatants were loaded into polyacrylamide gel.

### Western Blotting

30 μg of total protein extract, or 10 uL of proteins isolated from the culture medium as described in the *Protein extraction* paragraph, were separated using a gradient (4-20%) sodium dodecyl sulfate-polyacrylamide gel (SDS-PAGE) and transferred onto a polyvinylidene difluoride (PVDF) membrane as already described by Pecce et al. in 2023 ^27^. After 2 h of blocking using not-fat dry milk at 5%, proteins were detected with specific primary antibodies at a dilution of 1:2000: ANGPTL3 (GeneTex), PCSK9 (Abcam), a-ACTIN (Sigma Aldrich), Albumin (Abcam) incubated overnight at 4°C. After the incubation of horseradish peroxidase (HRP)-conjugated secondary antibodies, the bands were detected with chemiluminescence using Clarity Western ECL substrate (BIO-RAD) and a charge-coupled-device camera (Chemidoc, BIO-RAD). Alternatively, the bands were detected with a colorimetric precipitate using alkaline-phosphatase (AP)-conjugated secondary antibodies and 5-bromo-4-chloro-3-indolyl-phosphate/nitro blue tetrazolium substrates (Promega).

### Co-immunoprecipitation

Co-immunoprecipitations were performed using Dynabeads G (Thermofisher Scientific). 50 µL of Dynabeads were neutralized with a wash in Phosphate Buffer Saline (PBS) solution and then binded with 1 µg of specific antibody diluted 1:100 in PBS. Beads were then washed twice and resuspended in 100 µg of total protein extract in 1 mL of PBS in overnight gentle rotation at 4°C. Beads were then magnetically separated from total proteins, the antibody-bead complex was dissociated using a loading buffer, and beads were boiled in a heat bath at 95°C and loaded in SDS-polyacrylamide gel for electrophoretic separation as described in *Western Blotting* section.

### Immunofluorescence

Immunofluorescence was performed using fixed cells in 4% paraformaldehyde (PFA) for 10 minutes, and then the cells were washed twice in phosphate buffer solution (PBS). Cells were then permeabilized using PBS-Triton-X solution 0,01%, and blocking was performed using a solution of PBS-bovine serum albumin (BSA) 3% for 2 hours. Primary antibodies were then incubated overnight at a dilution of 1:50. The day after, secondary antibodies were incubated for 45 minutes, cells were washed twice, and nuclei were stained with the 4′,6-diamidino-2-phenylindole (DAPI) at a dilution of 1:5000. The fluorescence was conserved with a drop of mounting solution (1:1 PBS and glycerol) on samples and acquired using the THUNDER imaging system inverted microscope (Leica).

### Quantitative PCR

RNA was isolated from cells using RNeasy Mini Kit (Qiagen, Hilden, Germany) and quantified with Nanodrop 2000 (Thermo Fisher Scientific). The High-Capacity cDNA Reverse Transcription Kit (Thermo Fisher Scientific) synthesized cDNA from RNA according to the manufacturer’s instructions (Thermo Fisher Scientific). The expression levels of *ANGPTL3*, *PCSK9* and transcripts related to lipogenesis (*SREBP-1c, ACACA, ACLY* and *FASN*) and lipolysis (*PPAR-alpha* and *CPT1A*) were analyzed on cDNAs from treated and control HepG2 cells using TaqMan Gene Expression assays [CPT1A HS00912671; PPARα HS00947536; SREBP1 HS04984975; ACACA HS01046047; ACLY HS00982738; FASN HS01005622] and TaqMan Universal Master Mix according to the manufacturer’s instructions (Thermo Fisher Scientific). Results were calculated using the 2−ΔΔCt method and normalized using b-Actin as endogenous control (TaqMan® Gene Expression Assays HS99999903). Data were expressed as the mean ± standard deviation (SD) of three replicates relative to the CTRL sample that was used as a calibrator.

### Oil Red O cell staining

For Oil Red O staining, cells were fixed using a solution of PFA 0.75% for 5 minutes; cells were washed with PBS and then with isopropyl alcohol. Cells were stained with Oil Red working solution (40% Oil-Red O and 60% of isopropyl alcohol) for 5 minutes and washed with distilled water twice. For de-staining and Oil Red quantification, paraffin fixed cells were incubated with 1 mL of Isopropanol until complete destaining; solution absorbance was then read at 450 nm using the Varioskan™ plate reader (Thermo Scientific).

### Flow Cytometry

Cells were labeled with the fixable viability dye eFluor780 (Thermo Fisher Scientific) for 30 minutes at room temperature. Then, they were stained for surface markers for 20 min at 4°C in PBS 2% FBS with LDLR BV605 (BD 745174). All flow cytometry experiments acquired cells on LSR Fortessa (BD Biosciences). Data was analyzed using FlowJo software, version 10.0.8r1 (BD Biosciences).

### Oil Red O LDL staining

Bini et al., in 2024 ^28^, described this protocol for monitoring LDL uptake in the time-lapse experiment. In brief, 150 µg of LDLs were diluted in 100 µL of PBS, stained with 50 µL of Oil Red O reconstituted in DMSO, and centrifuged at 4000 RPM for 4 minutes. The obtained solution of Oil Red-stained LDLs at 1µg/µL was used for time-lapse LDL-uptake monitoring using the THUNDER imaging system inverted microscope (Leica).

### Statistical analysis

Statistical analyses were performed with GraphPad Prism 8.01. Differences between continuous variables were assessed with the unpaired t-test for cell lines. Two-way ANOVA was used to analyze flow cytometry data. A p-value < 0.05 was considered statistically significant.

## RESULTS

### ANGPTL3 and PCSK9 show intracellular and extracellular interaction in HepG2 cells

Based on the observation of reduced levels of circulating PCSK9 in patients carrying the homozygous LOF in ANGPTL3, we hypothesized the existence of a direct interaction between ANGPTL3 and PCSK9 proteins. Firstly, we used AphaFold2 software to test this hypothesis by modeling a potential molecular docking into a complex. We used the full-length sequences of the two proteins and their mature forms as input for the computation and studied the different hypothetical interactions: 1) full-length PCSK9 and ANGPTL3, 2) mature PCSK9 and ANGPTL3 C-terminus domain, 3) mature PCSK9 and ANGPTL3 N-terminus domain, 4) full-length ANGPTL3 and mature PCSK9. As shown in **Figure S1**, 3 out of the 4 hypothetical complexes showed high spatial uncertainty, leading us to deduce that these complexes do not exist. In particular, in the full-length/full-length complex, we obtained a low pLDDT score, and we found that the areas showing higher uncertainty were the PCSK9 pro-peptide (known to be cleaved and folded in the putative PCSK9 active site^14,15^) and ANGPTL3 N-terminus domain. The predicted complexes using the N-terminus and C-terminus domains of ANGPTL3 show high folding scores. However, they interact with PCSK9 in sites different from the full-length ANGPTL3 and small areas, making an unstable interaction. As reported in **Figure 1**, the highest score of pLDDT was obtained with ANGPTL3 full-length and mature PCSK9. This potential direct interaction shows the highest odds of existence. This computed model shows a pLDDT of 72.9, a pTM of 0.61, and an ipTM of 0.57. In this predicted complex, ANGPTL3 docks with PCSK9 in several aminoacidic residues (Thr347-Leu351; Asn425-Phe429; Ser662-Glu669), thus covering some residues that are crucial for PCSK9 interaction with LDL receptor (LDLR) such as Asp374 and Ser127^29^. Indeed, in our model, both sites are covered by ANGPTL3 residues that maintain a distance of around 20Å, which is sufficient to determine steric hindrance^30^ (**Figure 1B**-**1C**). These computed data confirm the potential interaction between ANGPTL3 and mature PCSK9, suggesting that ANGPTL3 might determine a steric hindrance in PCSK9.

**Figure 1.**
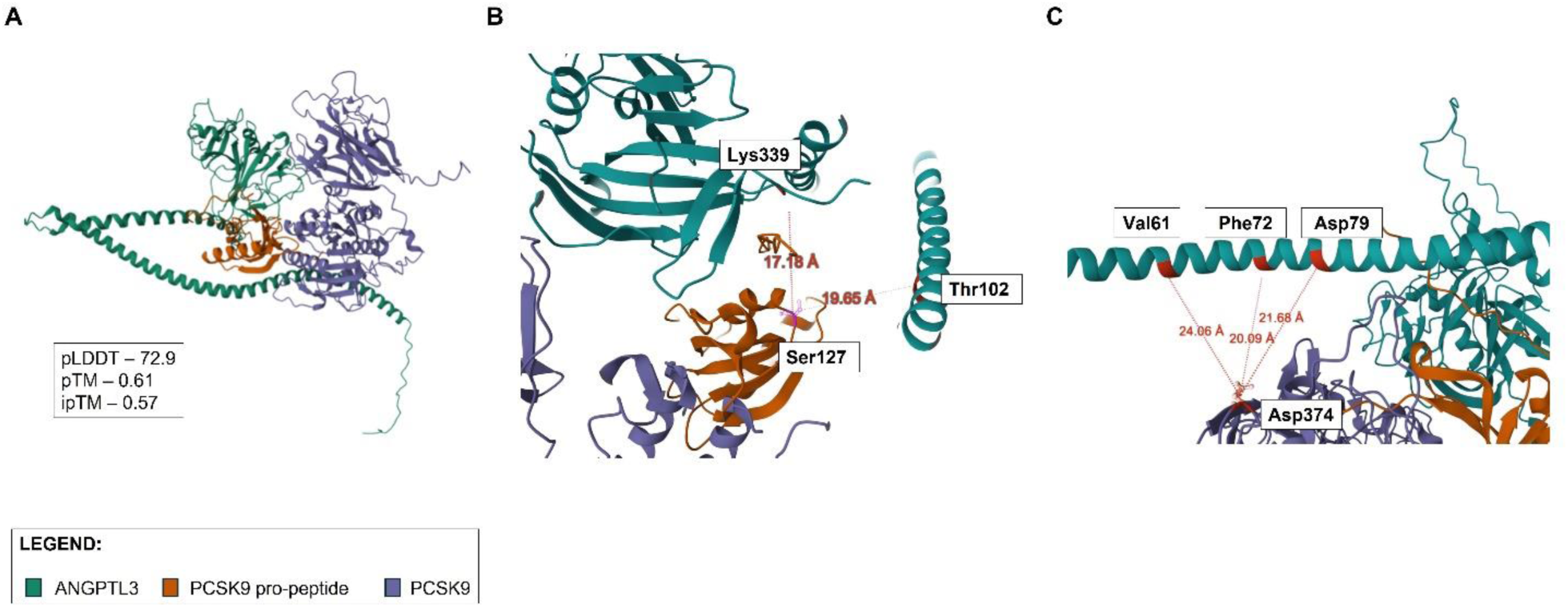
In silico prediction of ANGPTL3-PCSK9 interaction. **Panel A** shows the computed interaction between full-length ANGPTL3 and mature PCSK9 (PCSK9 propeptide was inserted as a different folding peptide). **Panel B** shows ANGPTL3 that covers Ser127 residue in PCSK9 pro-peptide. **Panel C** shows PCSK9 Asp374 residue covered by ANGPTL3 N-terminus domain. A pLDDT>70 is considered accurate, with a pTM>0.5 and an ipTM>0.5.

To verify the interaction predicted *in silico*, we performed co-immunoprecipitation (Co-Ip) experiments. Since ANGPTL3 and PCSK9 are secreted proteins, we first verified protein-protein interaction in the human hepatic cell line HepG2 culture media. In the fraction immunoprecipitated with the antibody for ANGPTL3 (IP-ANGPTL3), we found the mature form of PCSK9 (mPCSK9). In the same way, in the fraction immunoprecipitated with the antibody for PCSK9 (IP-PCSK9), we found ANGPTL3 (**Figure 2A**). These results are compatible with a physical interaction between ANGPTL3 and PCSK9. Then, we evaluated if the same interaction could be found at the intracellular level. To this aim, we used the total protein extract of HepG2 cells and repeated the Co-Ip experiment. Also, in this case, in the IP-ANGPTL3 sample, we identified PCSK9, and conversely, in the IP-PCSK9 sample, we found ANGPTL3 (**Figure 2B**). Then, we performed immunofluorescence to confirm this data. We used the HepG2 cells to co-localize the two proteins and confirm the intracellular interaction. As shown in **Figure 2C**, the signals corresponding to ANGPTL3 (green) and PCSK9 (red) co-localize in the cytoplasm of the cells near the nuclear membrane. These findings confirm that ANGPTL3-PCSK9 tends to form a complex, highlighting that the interaction is maintained intracellularly and extracellularly.

**Figure 2.**
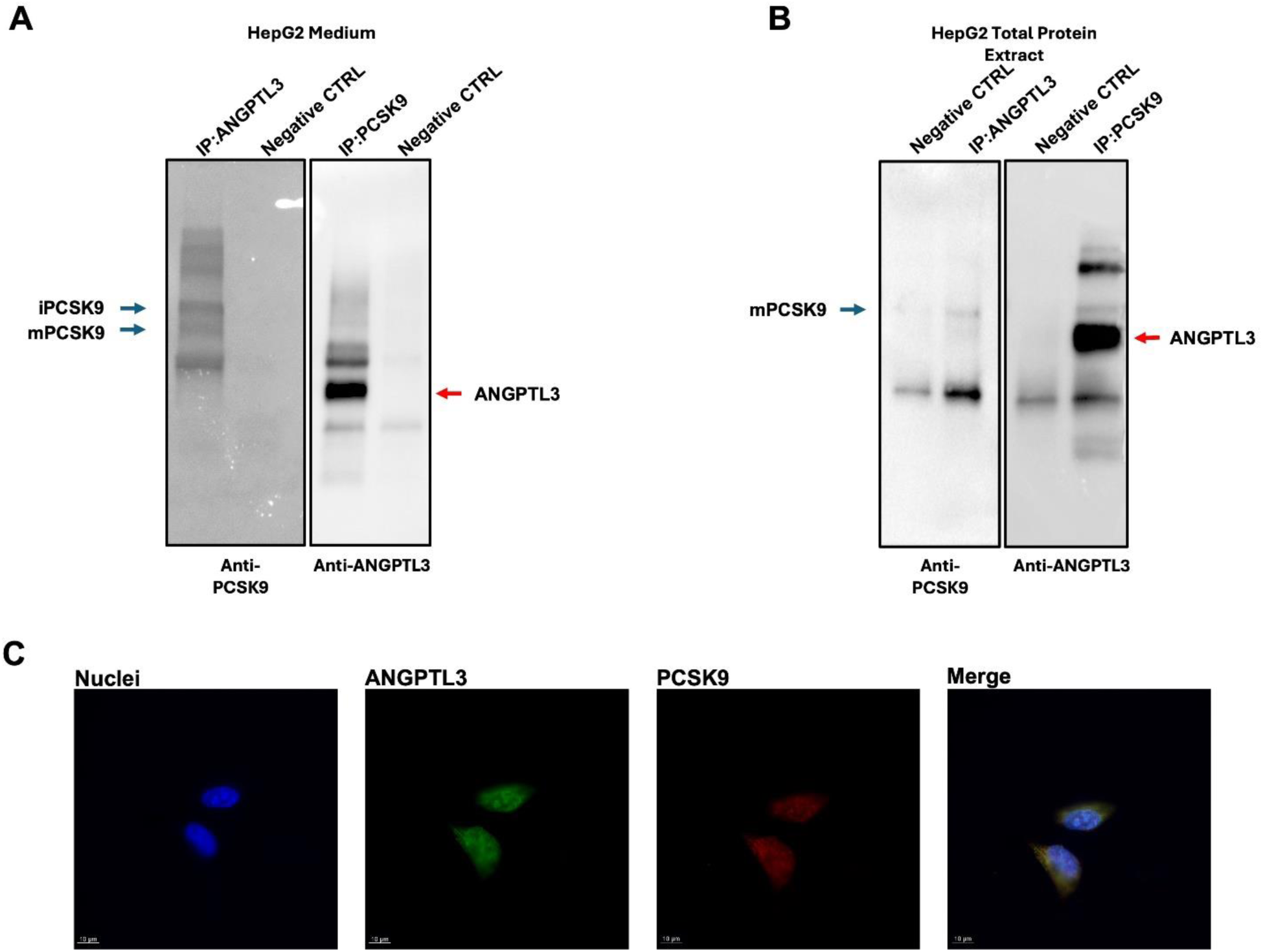
The interaction between ANGPTL3 and PCSK9. **Panel A** shows the Co-Immunoprecipitation (Co-IP) of total proteins isolated from the growth medium. IP:ANGPTL3 and IP:PCSK9 samples contain proteins isolated from 2 mL of growth medium using beads bound with 2µg of ANGPTL3 or PCSK9 antibody, respectively. Negative CTRL contains proteins isolated from a 2 mL medium using unbound beads. **Panel B** represents Co-IP of 100 ug of total protein extract from HepG2 cells. IP:ANGPTL3 and IP:PCSK9 contain protein isolated from beads bound with 2µg of PCSK9 or ANGPTL3 antibody. Negative CTRL represents proteins from unbound beads. Anti-PCSK9 and anti-ANGPTL3 represent the antibodies hybridized on the membrane. **Panel C** shows the immunofluorescence of HepG2 cells formalin-fixed. Cell nuclei are stained with DAPI and shown in blue. ANGPTL3 is shown in green. PCSK9 is shown in red. The merged red-green image shows co-localization as yellow spots. The acquisitions are performed at 63X magnification. The scale bar is 10 µm as a reference. iPCSK9 = immature PCSK9; mPCSK9 = mature PCSK9; ANGPTL3 = full-length ANGPTL3.

### Nutrients availability modulate ANGPTL3 and PCSK9 interaction

After establishing that the ANGPTL3 and PCSK9 proteins can interact with each other, we searched for factors that can influence this interaction. To this aim, we first generated HepG2 cell lines overexpressing *ANGPTL3* (oeANGPTL3), overexpressing *PCSK9* (oePCSK9), or both genes (oeANGPTL3-PCSK9) to enhance the activity involving one, the other, or both proteins. Consistently, all overexpressing cells showed higher levels of ANGPTL3 and PCSK9 protein than control cells. In particular, oeANGPTL3 showed the highest ANGPTL3 protein level in the medium, while oePCSK9 cells showed the highest PCSK9 levels intracellularly. (**Figure S2**).

Then, we decided to analyze the levels of ANGPTL3 and PCSK9 in *feeding* and *fasting* conditions. To simulate these conditions, we used nutrient-rich or nutrient-poor medium, based on DMEM or F12 medium, respectively. The composition of the two media is reported in the **Supplementary Table 1**. In particular, to simulate *feeding*, we grew cells in a nutrient-poor medium, and after 48h, we switched to a nutrient-rich medium for 4h; conversely, the switch from a nutrient-rich to a nutrient-poor medium was used to simulate *fasting* (**Figure S3**).

Afterward, we analyzed differences in the expression levels of *ANGPTL3* and *PCSK9* transcripts in the different experimental conditions using real-time PCR (**Figure S4**). Our results indicated that, although nutritional change is a sufficient stimulus to regulate *ANGPTL3* and *PCSK9* transcript expression, all overexpressing cells maintain higher levels of *ANGPTL3* and *PCSK9* than those found in control cells. Overall, we conclude that the proposed overexpression system is efficient, and nutrients are effective modulators of transcript levels in overexpressing cells.

Once we established that our overexpressing system was functional, we analyzed ANGPTL3 and PCSK9 protein expression intracellularly and in the culture media (**Figure 3**). In control (CTRL) cells, ANGPTL3 levels increase in the feeding condition both intracellularly and in the culture medium. Conversely, in the fasting condition, ANGPTL3 is reduced in the culture medium, while ANGPTL3 levels do not change intracellularly. PCSK9 levels did not change in the culture medium of control cells in *feeding* and *fasting* conditions. Intracellularly, PCSK9 levels were reduced in fasting conditions, and its intracellular levels did not change in feeding. In oeANGPTL3 cells, ANGPTL3 protein levels are predominantly increased in the culture medium, whereas PCSK9 levels did not change compared to CTRL cells. In PCSK9oe cells, ANGPTL3 levels consistently increase intracellularly compared to CTRL and oeANGPTL3 cells. Conversely, PCSK9 levels are increased intracellularly and extracellularly in *feeding* and *fasting* conditions. In oeANGPTL3-PCSK9 cells, ANGPTL3 and PCSK9 levels change accordingly to CTRL cells intracellularly and in the control medium in the different nutritional conditions.

**Figure 3.**
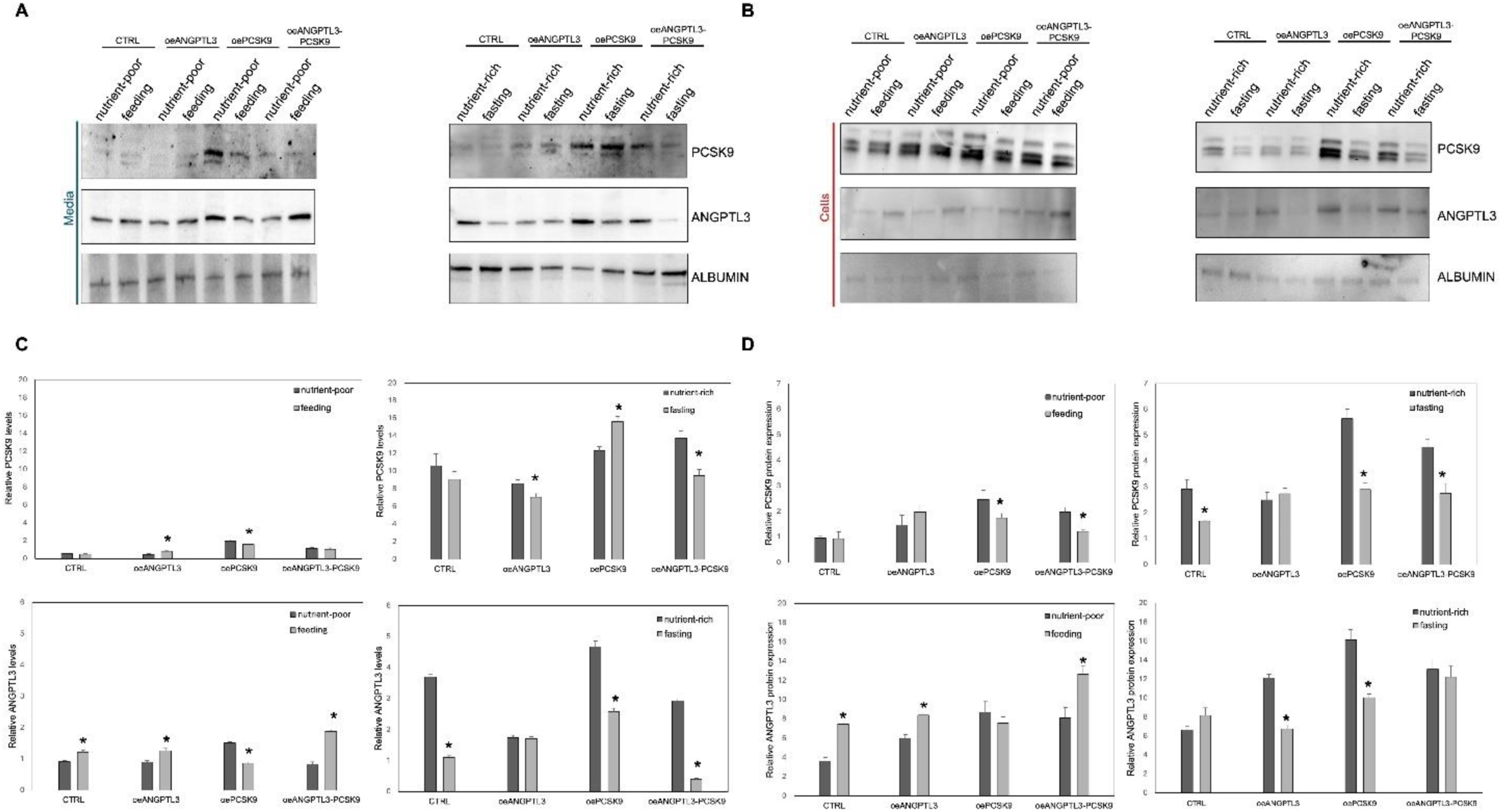
Nutrients modulate ANGPTL3 and PCSK9 levels - oePCSK9 increases ANGPTL3 expression. **Panel A** shows the representative western blot of the culture media. Proteins were extracted from 2 mL of FBS-free medium of cells growing in nutrient-poor or nutrient-rich medium, and simulating feeding condition (switching for 4h from nutrient-poor to nutrient-rich medium), or fasting condition (switching for 4h from nutrient-rich to nutrient-poor medium). **Panel B** shows the representative western blot analysis of 30 µg of total protein extract from cell lysate growth in the nutrient-poor or -rich medium and in *feeding* and *fasting* conditions. **Panels C** and **D** represent the densitometric analysis of three independent experiments. Data are shown as relative levels (for the media) and expression (for the cells) normalized for the quantity of the albumin. Asterisks indicate statistically significant change (*p*<0.05), test applied: Student T-test.

Overall, oePCSK9 cells show the highest intracellular levels of ANGPTL3 in nutrient-rich conditions, while oeANGPTL3 cells show the lowest intracellular levels in *fasting* conditions. These results suggest that PCSK9 may also regulate ANGPTL3.

Given the above-reported findings, we further investigated the behavior of the ANGPTL3-PCSK9 complex in *feeding* and *fasting* conditions. We performed co-immunoprecipitations to isolate the secreted forms of PCSK9 in the culture media of HepG2 cells and hybridized the anti-ANGPTL3 antibody on the western blotting membrane. As shown in **Figure S5A**, in nutrient-poor conditions, we observed the presence of ANGPTL3 in the samples immunoprecipitated with PCSK9 antibody (IP:PCSK9). Moreover, in the *feeding* starting sample (INPUT), ANGPTL3 appears in a great band under the 48 kDa (green arrow), indicating that the protein is cleaved in this condition, with the absence of bands corresponding to ANGPTL3 in the IP:PCSK9 sample. In **Figure S5B**, we observed the presence of the ANGPTL3-PCSK9 complex both in nutrient-rich and fasting conditions (red arrow indicates ANGPTL3 presence in PCSK9 precipitate). Conversely, we observed the presence of the ANGPTL3-PCSK9 interaction in the nutrient-poor culture media and the absence of the complex in *feeding* conditions. We can conclude that the ANGPTL3-PCSK9 complex exists in *fasting* conditions but not under *feeding* conditions.

These results suggest that nutrients modulate PCSK9 and ANGPTL3 at the post-translational level and, to a lesser extent, at the transcriptional level. The two proteins show physical interaction, but the complex is dissociated in the presence of nutrients. In other words, the ANGPTL3-PCSK9 complex exists at the intracellular level in fasting conditions, while in feeding conditions, ANGPTL3 and PCSK9 are released, and they function as unconjugated proteins.

### The LDL treatment releases PCSK9 and ANGPTL3 from the complex

Once we observed the modulation of the ANGPTL3-PCSK9 complex in response to nutrients in lipid-free media, we challenged our system with LDLs to study the effect of this lipoprotein on the complex’s modulation. As shown in **Figure 4**, in the absence of LDL, we confirmed the intracellular co-localization of ANGPTL3 and PCSK9 in CTRL cells as well as in cells overexpressing *ANGPTL3*, *PCSK9*, or both genes. In the presence of LDL, both CTRL and oeANGPTL3-PCSK9 cells maintain part of the ANGPTL3 and PCSK9 co-localization. Conversely, the complex seems to dissociate entirely in oeANGPTL3 and oePCSK9 cells. This latter observation can be explained by hypothesizing that the presence of LDL binding to the specific cellular receptor can generate the dissociation of the ANGPTL3-PCSK9 complex like the increase in non-lipid nutrients in the growth medium.

**Figure 4.**
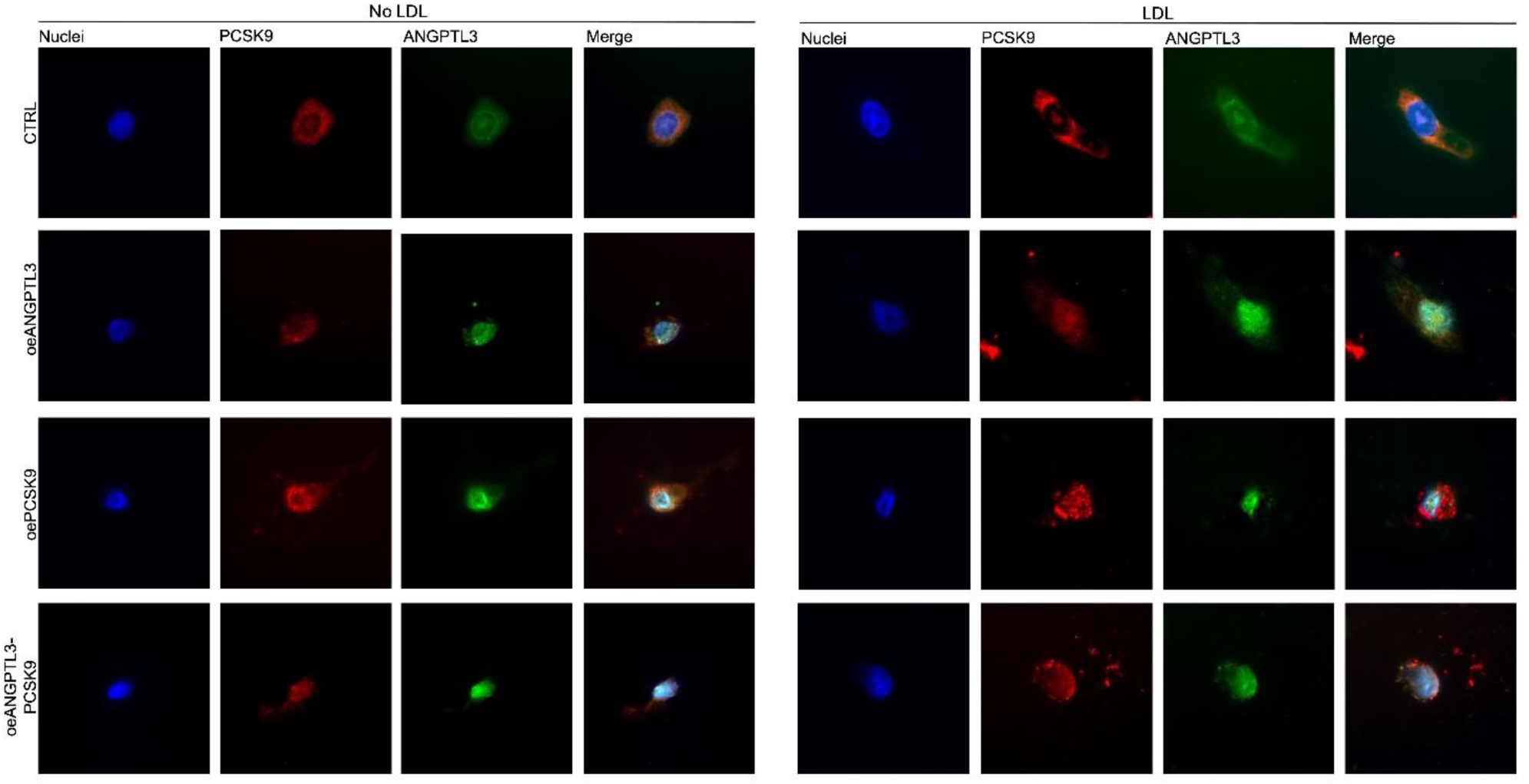
ANGPTL3 and PCSK9 complex dissociates in the presence of LDL. Immunofluorescence of HepG2 cells treated or not with 10 µg of LDL-C for 4 hours. Cell nuclei are shown in blue, PCSK9 in red and ANGPTL3 in green. ANGPTL3-PCSK9 co-localization is represented in yellow in the merged figure.

### ANGPTL3 is involved in ApoB maturation, while PCSK9 determines ApoB degradation

Subsequently, we investigated the role of *ANGPTL3* and *PCSK9* overexpression in modulating lipoprotein synthesis and secretion. In particular, we analyzed the levels of ApoB accumulation in the culture media and intracellularly in nutrient-rich and nutrient-poor medium for 24h at 5 different time points. We started the analysis after 48h from overexpression of *ANGPTL3*, *PCSK9,* or both genes in HepG2 cells (T0). Then, after refreshing the medium, we analyzed the levels of the secreted and the intracellular ApoB after 4h, 16h, 20h, and 24h (**Figure 5**). In nutrient-rich, the control cells reached the peak of ApoB concentration in the media after 16h. We observed that at the same time point, the intracellular levels of ApoB reached the lowest level, thus suggesting that the protein produced is all secreted. In the same conditions, the accumulation of ApoB in the oeANGPTL3 cells was maintained constant over 24h. In the oePCSK9 cells, ApoB accumulation reached the highest levels; on the contrary, in the oeANGPTL3-PCSK9 cells, ApoB showed the lowest levels of accumulation (**Figure 5A-B-C-D**). At the intracellular level, we observed different bands corresponding to ApoB isoforms. If compared with control cells, the oeANGPTL3 cells showed an upper band (molecular weight MW > 100 KDa), representing ApoB with post-translational modifications (e.g., glycated isoform of ApoB)^31^. On the contrary, the oePCSK9 and oeANGPTL3-PCSK9 cells showed lower bands (MW < 100 KDa), representing ApoB catabolic fragments^31^ (**Figure 5C**). In the nutrient-poor medium, the peak of ApoB secretion in CTRL cells is observed at 20h. The oeANGPTL3 anticipated the maximum secretion peak at 4h with a second peak at 20h. The oePCSK9 cells also showed higher levels of ApoB accumulation in this condition, with a higher peak at 4h. The oeANGPTL3-PCSK9 cells showed the lowest levels of ApoB accumulation during the 24h. ApoB fragments are equally represented in the four experimental conditions at the intracellular level. These results allow us to conclude that the simulated condition of *feeding* and *fasting* can regulate ApoB synthesis, maturation, and secretion. Indeed, in a nutrient-poor medium, we observed a slower peak of accumulation of ApoB and lower intracellular and extracellular ApoB levels in CTRL cells. When *ANGPTL3* is overexpressed, the maturation of ApoB is stimulated, as supported by the evidence of intracellular accumulation of higher molecular weight ApoB. The overexpression of *PCSK9* accelerates the degradation of ApoB, leading to higher amounts of intracellular bands and catabolic fragments of the protein. The overexpression of both genes led to a faster turnover of ApoB.

**Figure 5.**
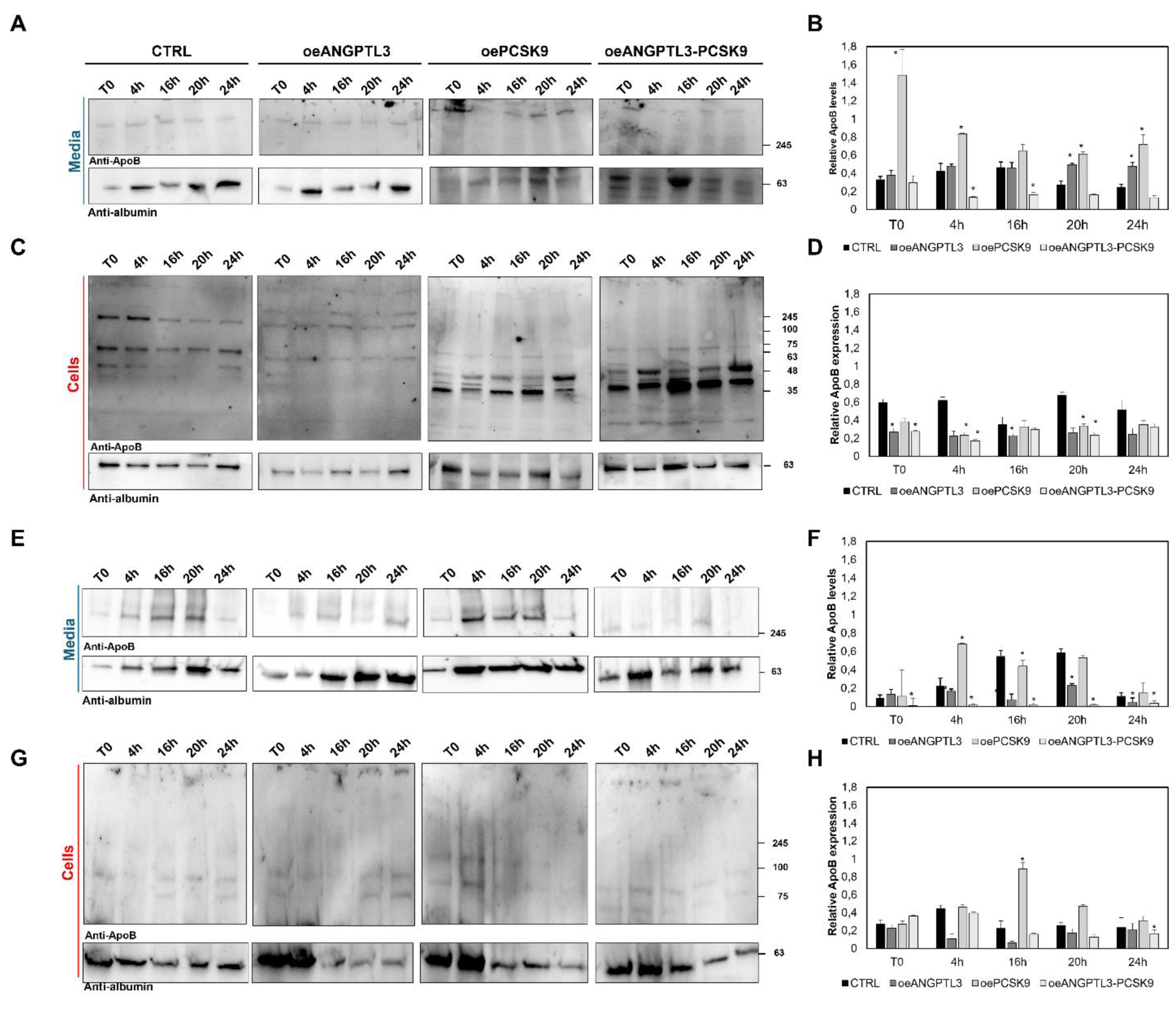
The overexpression of ANGPTL3 accelerates the maturation of ApoB, and the overexpression of PCSK9 accelerates its degradation. Secreted and intracellular levels of ApoB throughout 24h at five different time points. Representative western blot (right panels) and densitometric analysis (left panels) of ApoB levels in HepG2 cell line (CTRL) overexpressing ANGPTL3 (oeANGPTL3), PCSK9 (oePCSK9) or both genes (oeANGPTL3-PCSK9). Cells were grown in nutrient-rich conditions, and ApoB was measured in media (panels A-B) and in the cells (panels C-D). Cells were grown in nutrient-poor conditions, and ApoB was measured in the media (panel E-F) and in the cells (panel G-H). The densitometric analysis only considered bands migrating at a predicted MW> 100 KDa. That corresponds to nascent and maturing ApoB. Asterisks indicate statistically significant change (*p*<0.05), test applied: Student T-test.

Given that literature data highlight the role of PCSK9 in recycling LDLR, we further decided to investigate LDL receptor membrane levels using the flow cytometry technique. Results are shown in **Figure S6**. Membrane LDLR expression is maximal in the nutrient-poor condition and consistently reduces in *feeding*. Minimal LDLR expression is observed in the nutrient-rich condition, and membrane expression is increased in *fasting*. No significant differences were observed between the different overexpressing conditions and control cells. We can conclude that LDLR expression is predominantly regulated by nutrients in our growth condition.

### Intracellular lipid accumulation is determined by relative levels of ANGPTL3 and PCSK9 and is independent from the *de novo* lipogenesis

After observing the effects of the overexpression of *ANGPTL3* and *PCSK9* on ApoB metabolism, we analyzed the effects of these proteins on the intracellular storage of neutral lipids by using Oil Red-O staining in cells overexpressing *ANGPTL3, PCSK9*, or both genes in nutrient-rich or nutrient-poor culture conditions. This set of experiments was based on the rationale that the synthesis, degradation, and secretion of apoB are crucial determinants of neutral lipid accumulation in the liver cells^32^.

As shown in **Figure 6A-B**, overall lipid accumulation in nutrient-rich medium appears to be more organized in intracellular vesicles, while lipid accumulation in nutrient-poor medium appears more interspersed. In nutrient-rich conditions, oeANGPTL3 cells show a 15% reduction in intracellular lipid accumulation compared with CTRL, while oePCSK9 and oeANGPTL3-PCSK9 cells show a decrease of 20% and 40%, respectively.

**Figure 6.**
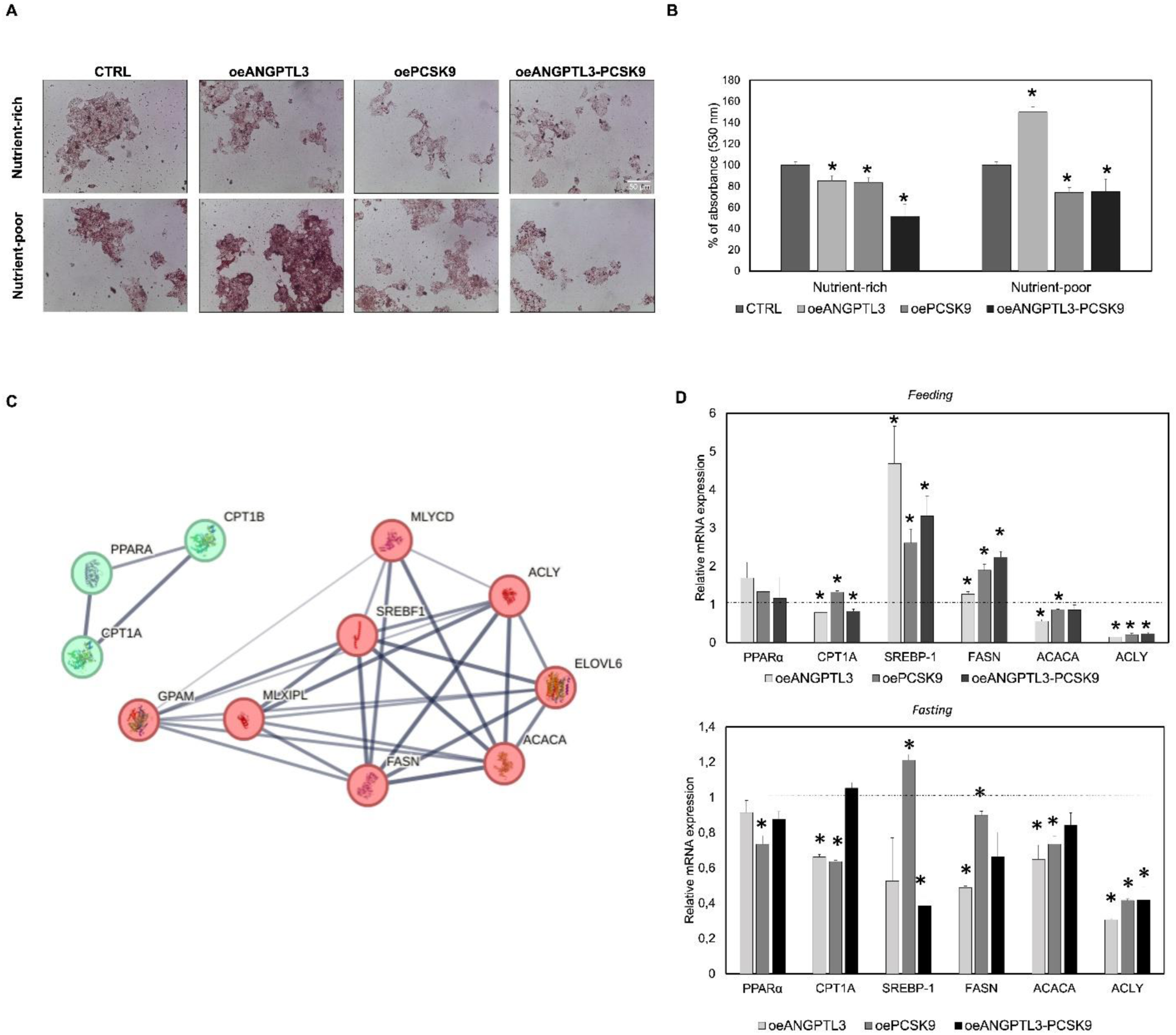
*ANGPTL3* increases intracellular lipid levels in nutrient-poor growth conditions independent from *de novo* lipogenesis, and the overexpression of *PCSK9* reduces intracellular lipid content associated with a reduction of *de novo* lipogenesis. **Panel A** shows the Oil Red-O staining of HepG2 control cells or overexpressing ANGPTL3, PCSK9, or both genes, growth in the nutrient-rich or nutrient-poor medium. Panel B shows the Oil Red absorbance at 520 nm wavelength of the dye extracted from the stained slide, expressed as a percentage relative to control. **Panel C** shows STRING-based protein-protein interaction analysis in beta-oxidation pathways (green) and lipogenesis (red). **Panel D** shows mRNA expression levels for different lipogenesis or lipolysis genes in overexpressing HepG2 cells in *feeding* and *fasting* conditions. Relative mRNA expression is normalized to control cells (dotted line). Asterisks indicate statistically significant change (*p*<0.05), test applied: Student T-test.

In nutrient-poor conditions, oeANGPTL3 cells show a 50% increase in lipid accumulation compared with control cells; conversely, the overexpression of *PCSK9* and both genes show a 20% reduction in lipid accumulation. As shown in **Figure 3**, the highest levels of intracellular ANGPTL3 and PCSK9 are observed in nutrient-rich conditions. On the other hand, in the nutrient-poor medium in oeANGPTL3, we observed lower levels of both proteins. These results led us to conclude that there is lipid intracellular accumulation if ANGPTL3 and PCSK9 are expressed at lower levels and a reduced lipid accumulation when ANGPTL3 and PCSK9 are expressed at higher levels.

To better explain the observed differences in lipid accumulation, we analyzed the expression levels of selected targets regulating lipolysis and lipogenesis in all overexpression cells under different growth conditions. As shown in the STRING model reported in **Figure 6C**, we selected targets of lipogenesis clustered around *SREBP-1c* (*SREBP-1c, ACACA, ACLY*, and *FASN*) and targets of enhancing beta-oxidation clustered around *PPARα* (*PPARα* and *CPT1A*). As reported in **Figure 6D**, in *feeding* conditions, cells overexpressing *ANGPTL3* show a reduction in the expression of *CPT1a*, indicating a reduced activation of mitochondria beta-oxidation and a 5-fold increase in the expression of *SREBP1c* with a decreased expression of *ACLY*, thus determining a reduction in lipogenesis. The oePCSK9 cells show an increased expression of *CPT1a* with a decreased expression of *ACLY*, therefore meaning an increase in beta-oxidation and a reduced lipogenesis; cells overexpressing both proteins also show slightly reduced expression of *CPT1a* and *ACLY*. Surprisingly, in all overexpression conditions, the increased expression of *SREBP-1c* is not associated with the increase in *ACLY* or *ACACA* levels^33,34^, suggesting that the changes observed in lipid levels do not involve *de novo* lipogenesis. As shown in **Figure 6D**, in *fasting* conditions, the overexpression of *ANGPTL3* determines a reduction in the expression of *CPT1a* and *ACLY*, as observed in the *feeding* condition. Differently from *feeding*, in *fasting* conditions, the expression of *SREBP-1c* is comparable to the control cells. In cells overexpressing *PCSK9*, the expression of *CPT1a* is reduced compared to the feeding condition, while the expression of *ACLY* is reduced, and the one of *SREBP-1c* is increased like in *feeding*. In cells overexpressing both genes, growth in *fasting* conditions, mRNA levels of *SREBP-1c* are reduced, and levels of *CPT1a* are comparable to controls. As observed in *feeding* conditions, *ACLY* and *ACACA* levels are reduced compared to the control. These results suggest that the overexpression of *ANGPTL3* increases intracellular lipid levels in nutrient-poor growth conditions independent from *de novo* lipogenesis. In addition, the overexpression of *PCSK9* reduces intracellular lipid content, which is associated with a reduction of *de novo* lipogenesis.

### ANGPTL3 induces LDL catabolism while PCSK9 determines intracellular LDL entrapment

In the previous results, we have observed that ANGPTL3 and PCSK9 determined a similar reduction in intracellular lipid accumulation in nutrient-rich conditions. We also observed that LDL treatment determines the dissociation of the ANGPTL3-PCSK9 complex. We treated the cells with LDLs isolated from a pooled plasma of healthy donors stained with Oil Red-O. Then, we observed the LDL uptake in the control and overexpressing samples using time-lapse and monitoring cells for 20h.

As shown in **Video S1,** CTRL cells internalize the stained LDLs (seen as black dots), which disappear over time. In the oeANGPTL3 cells, we observed a faster disappearance rate of black dots compared with CTRL cells, and eventually, some cells exploded at the end of the observation. The oePCSK9 cells show a higher quantity of black dots internalized over time that can be observed for all 20h of the observation. The oeANGPTL3-PCSK9 cells showed similar behavior to CTRL cells.

Moreover, we quantified the total internalized LDL, analyzing the red vesicles found in the cells in all acquisition frames using the ImageJ software; data were expressed as a total red area. As shown in **Figure 7A**, the overexpression of *ANGPTL3* determined a global reduction of the total red area, suggesting a decrease in the number of intracellular LDLs compared with the control sample. On the other hand, the overexpression of *PCSK9* determines an increase in the total red area if compared with those found in the control sample. Cells overexpressing both *ANGPTL3* and *PCSK9* showed a progressive increase of the intracellular LDL levels, which peaked at 11 hours and returned to levels comparable to those observed in the oeANGPTL3 sample.

**Figure 7.**
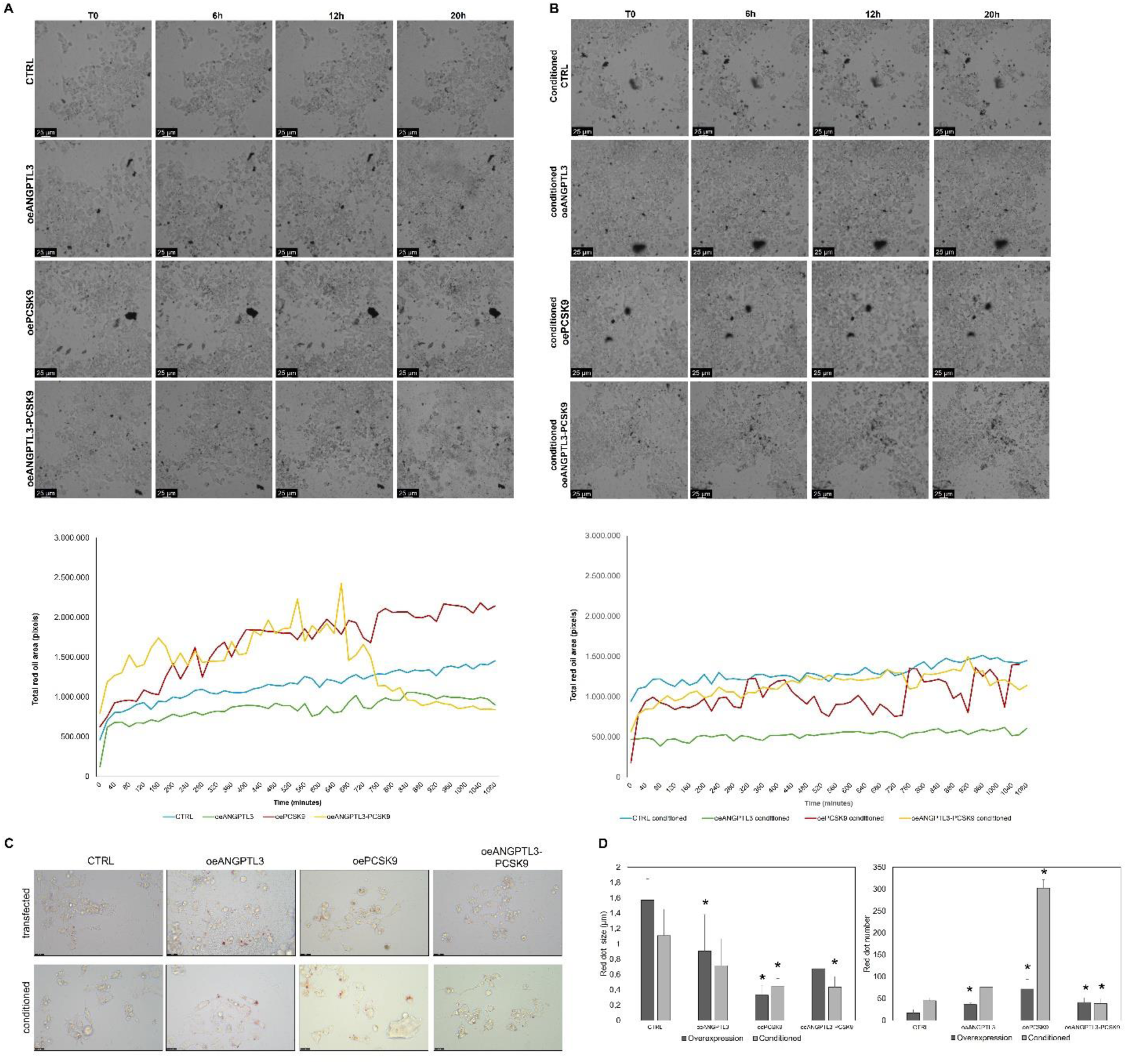
The oePCSK9 increases intracellular LDL levels, and oeANGPTL3 increases LDL turnover in HepG2 cells. (**A**) Representative frames of timelapse experiment with quantifying red area over time for HepG2 CTRL cell or overexpressing ANGPTL3 or PCSK9 or both. (**B**) Representative frames of timelapse of HepG2 cells conditioned with the medium of oeANGPTL3, oePCSK9, or oeANGPTL3-PCSK9 overexpressing cells. The quantification of the red area is represented in the bottom panel. (**C**) representative microphotographs of HepG2 cells 24h after the LDL treatment and (**D**) quantification of the number and dimension of the lipidic drop in transfected or conditioned cells. Asterisks indicate significative change if compared to the respective CTRL condition. Asterisks indicate statistically significant change (*p*<0.05), test applied: Student T-test.

We repeated the experiments using HepG2 cells pretreated for 24h with the medium obtained from oeANGPTL3, oePCSK9, and oeANGPTL3-PCSK9 samples (**Figure 7B**). These conditioned media allowed us to investigate the effect of secreted ANGPTL3 and PCSK9 on LDL internalization. As previously observed in the oeANGPTL3 sample, oeANGPTL3-conditioned cells also showed a reduction in the intracellular LDL, resulting from an analysis of the total red area. In the oePCSK9 conditioned cells, the levels of intracellular LDLs fluctuate over the 20h of observation, remaining at levels of the total red area as observed in the CTRL sample. In the oeANGPTL3-PCSK9 conditioned cells, the total red area observed was restored to those seen in control cells.

These results allow us to conclude that ANGPTL3 functions extracellularly as a signal to induce LDL catabolism. Indeed, the effects of ANGPTL3 overexpression are observed also in the conditioned medium. Conversely, PCSK9 determines the intracellular entrapment of LDLs, determining an increase in the measured total red area.

We also analyzed intracellular vesicle size at the end of the time-lapse experiment. Control cells show a small number of red vesicles of bigger size (> 1µm^2^), while cells overexpressing *ANGPTL3*, *PCSK9*, or both genes and cells treated with conditioned medium show a high number of smaller vesicles. Conditioned oePCSK9 cells have a number of vesicles per cell > 100, with a size < 0.4 µm^2^, **Figure 7C-D**. Supporting the observation of an LDL entrapment when *PCSK9* is overexpressed.

To verify the effects of ANGPTL3 and PCSK9 on LDL, we reproduced the experiment in two different non-liver-derived immortalized cell lines: HEK293 and HeLa. As shown in **Figure S7**, the overexpression of *ANGPTL3* also maintains a lower intracellular quantity of LDL in non-liver cells. On the contrary, the effect of the overexpression of *PCSK9* in non-liver cells is more evident using the conditioned medium; the cells treated with medium from oePCSK9 cells showed higher levels of the red area if compared with the control. Different from HepG2 cells, the HEK293 and HeLa cells tend to form smaller red vesicles of ̴ 0.8 µm^2^ for HeLa and ̴ 0.5 µm^2^ for HEK293, **Figure S7**.

When PCSK9 is overexpressed in all the investigated cell lines, endocytic vesicles are kept near the membrane, and bigger secondary endosomes appear much later. In contrast, ANGPTL3 seems to enhance the catabolism of LDL particles in this condition; endocytic vesicles rapidly disappear into secondary endosomes.

## DISCUSSION

Our findings provide several new insights into the intracellular hepatic role of ANGPTL3 and PCSK9. We demonstrated that ANGPTL3 and PCSK9 can form a complex that can be detected both intracellularly and in the cellular culture medium of HepG2 cells. We have also proved that this complex is dynamically regulated by nutrient availability as it forms in nutrient shortage conditions and dissociates in nutrient abundance conditions. LDL treatment is a sufficient stimulus to induce the dissociation of the ANGPTL3-PCSK9 complex. The two free proteins also regulate intracellular lipid and lipoprotein metabolism. In particular, ANGPTL3 enhances ApoB maturation, whereas PCSK9 increases ApoB intracellular degradation. The two proteins also regulate intracellular lipid accumulation: the overexpression of *ANGPTL3* determines intracellular lipid accumulation in nutrient-poor growth conditions, whereas *PCSK9* overexpression determines a reduction in intracellular lipid accumulation. ANGPTL3 and PCSK9 have also been found to regulate the uptake of LDLs finely. In cells overexpressing *ANGPTL3,* LDLs show a reduced intracellular accumulation over time, suggesting a faster LDL metabolism. Conversely, cells overexpressing *PCSK9* show increased LDL accumulation over time, suggesting a reduced LDL metabolism and particle entrapment.

We also observed a coordinated expression of ANGPTL3 and PCSK9 proteins. This offers a key to interpreting the observation by *Fazio et al.*^18^ that FHBL2 patients show reduced levels of circulating PCSK9. If the overexpression of PCSK9 determines the increase in the intracellular levels of ANGPTL3, we might postulate that the absence of ANGPTL3 may directly lead to a reduction in PCSK9 levels. Further, the coordinated regulation of the levels of these two proteins may also offer a further interpretation of the hypocholesterolemic phenotype associated with ANGPTL3 deficiency^35^. It is well known that the PCSK9 protein represents a potent inhibitor of LDLR activity, as it reduces the availability of receptor protein on the cell surface ^36^. Then, the reduced concentration of PCSK9 observed in individuals lacking ANGPTL3 could increase the LDLR-dependent catabolism of LDL particles and other lipoproteins containing ApoB, thus determining the hypocholesterolemic phenotype associated with FHBL2 ^35^. This hypothesis received some support from an *in vivo* kinetic study of ApoB-containing lipoproteins during therapy with Evinacumab, a monoclonal antibody that inhibits the function of ANGPTL3^37^. This study showed that treated individuals increased their fractional catabolic rate of ApoB-containing lipoprotein, mainly remnants.

Our findings suggest that the ANGPTL3-PCSK9 complex might act as a sensor in the liver, actively dissociating and reforming in response to variations in the influx of metabolic substrates into the liver, thus protecting hepatocytes from metabolic overload. In this regard, it is noteworthy that LDL abundance also promotes complex dissociation.

This paper compares the roles of these proteins in intracellular lipid metabolism, focusing on two key steps: ApoB secretion, which drives lipid secretion via VLDL, and the accumulation of neutral lipids in cells. Addressing these aspects appears relevant not only because the data presented in the literature in ANGPTL3 and PCSK9 knockout *in vitro* hepatic model do not appear univocal ^38–41^ but also because it can help to understand the results observed during pharmacological inhibition of the synthesis of these two lipoproteins.

We found that overexpressing ANGPTL3 enhances ApoB maturation, whereas PCSK9 increases ApoB intracellular degradation, and this effect was maximized in the presence of increased nutritional substrate availability. These findings may represent the counterevidence of those studies in which the expression of *ANGPTL3* was inhibited using the CRISPR/Cas 9 technique for the inactivation of the gene, where ApoB secretion was reduced ^41^. Some authors reported that the downregulation of ANGPTL3 determined by siRNA treatment was associated with increased neutral lipid accumulation *in vitro* models ^38,39^. A similar phenomenon has also been observed *in vivo* in humans in a recent clinical trial where patients were exposed to Vupanorsen, an antisense oligonucleotide (ASO) that targets *ANGPTL3* gene showed that the reduction of circulating levels of ANGPTL3 was associated with a dose-dependent liver fat accumulation^42^. Under our experimental conditions, the overexpression of ANGPTL3 determined intracellular lipid accumulation only in nutrient-poor conditions. Conversely, in nutrient-rich conditions, the overexpression of this protein was associated with a reduction in the cellular lipid contents, thus making it plausible that in these conditions, the absence of ANGPTL3 might promote the enrichment of hepatocytes with neutral lipids. These observations might be interpreted in light of what was theorized by *Hegele*^43^, who suggested that the liver lipid accumulation during pharmacological silencing of ANGPTL3 highly depends on the whole-body metabolic status, making ANGPTL3 a regulator of lipid intake and export.

We also observed that the overexpression of PCSK9 reduces intracellular lipid content, which was associated with a reduced expression of genes involved in *de novo* lipogenesis. This observation is in line with previous evidence indicating that plasma levels of PCSK9 are associated with liver steatosis^44^.

In this study, we described the functions of the ANGPTL3-PCSK9 complex only in cellular models and in simulated conditions of *feeding* and *fasting*. The regulation and function of this complex were not investigated in human subjects or murine models. However, we clarified the intracellular functions of ANGPTL3 and PCSK9 and their complex to the extent appropriate in a cellular model; future studies *in vivo* in human or murine models might provide a better understanding of their extracellular functions. In conclusion, our paper demonstrates that ANGPTL3 and PCSK9 regulate each other and intracellular lipoprotein metabolism by forming a novel protein complex regulated by nutrient availability. In addition, the presence of ANGPTL3 or PCSK9 in circulating lipoproteins might be crucial in regulating the lipoprotein half-life or targeting specific tissues and must be further investigated.

## ACKNOWLEDGEMENTS

**SB** and **VP**: study design, data production, analysis, and interpretation. **APG** and **SP**: data production, analysis, and interpretation. **LD, IM, ADC, SC,** and **DT**: data analysis and interpretation. **MA**: study conceptualization and data interpretation.

